# Free-edge mechanics and hydrodynamic coupling shape membrane remodeling after pore opening

**DOI:** 10.64898/2026.11.20.768126

**Authors:** Han Zhou, Yi-Wei Chang, Yoichiro Mori

## Abstract

Membrane remodeling after pore opening occurs in electroporation, vesicle rupture, intracellular delivery, and pathogen escape from host cells. During this process, a pore may expand or close while the surrounding membrane curls at the rim, forms a neck, or everts. Using a sharp-interface framework for open membranes in Stokes flow, we study how free-edge elasticity and the dissipation mechanism determine post-opening remodeling. The model combines Helfrich bending, Gaussian-curvature and line energies, and local area inextensibility with viscous dissipation in the membrane and the surrounding fluid. We compare it with a local-drag model driven by the same energy over a range of line tensions, Gaussian rigidities, and spontaneous curvatures. Starting from a spherical cap, the membrane first adjusts near the rim as it relaxes toward the free-edge moment condition. The subsequent pore dynamics depend strongly on line tension, with trajectories that either continue to expand or reverse and shrink. Gaussian rigidity changes the rim orientation; for positive Gaussian rigidity, a pronounced internal neck develops behind an outward-moving free edge. Increasingly negative spontaneous curvature localizes deformation near the rim and, at sufficiently negative values, causes the first complete rim turn to precede cap inversion. Hydrodynamic and local-drag simulations can dissipate similar amounts of energy while following different shape trajectories, so in the strongly curved regime the pathway depends on the dissipation model as well as on the energy. Together, the edge tangent angle, neck radius, cap depth, and the relative timing of rim rolling and cap inversion provide measurable geometric signatures that distinguish post-opening remodeling pathways beyond the pore radius.

## Introduction

During malaria parasite egress from red blood cells, the host membrane undergoes a sequence of large deformations: a pore opens, the pore rim curls outward, and the remaining membrane cap eventually everts as the parasite escapes [1, 2]. Similar post-rupture remodeling appears in electroporation, intracellular delivery, transiently ruptured lipid vesicles, and other cellular membrane-remodeling events [3–6]. Depending on the mechanical setting, the membrane may retract, retain a finite opening, form an internal neck, or roll near the free edge. Describing these pathways requires a model that resolves the pore edge together with the surrounding membrane shape.

Experiments on ruptured vesicles show that pore growth and closure depend on membrane tension, line tension at the pore edge, and fluid leakage [6, 7]. Molecular dynamics simulations resolve pore formation, lipid rearrangement near the edge, and the molecular origin of line tension [8–10], but at length and time scales smaller than those of whole-vesicle or whole-cell remodeling. At larger scales, reduced continuum models describe the dynamics in terms of pore radius, vesicle pressure, or a small number of geometric variables [11–13]. Such models capture important features of pore growth and resealing, but they do not resolve the coupled evolution of the free edge and the surrounding membrane geometry.

At the continuum scale, membrane shape is commonly described by Helfrich bending elasticity [14, 15], and the free edge of an open membrane carries a line energy that favors edge shortening. The Gaussian-curvature energy is constant for a closed membrane of fixed topology, but for an open surface it contributes directly to the force and moment conditions at the edge [16–18], and equilibrium calculations show that Gaussian rigidity can modify the membrane shape near a prescribed pore [19]. Spontaneous curvature provides an additional driving mechanism for rim curling and has been proposed in models of red-cell membranes and malaria parasite egress [2, 20]. These elastic forces are resisted by viscous dissipation within the membrane and in the surrounding fluid [21–23]. Diffuse-interface models naturally accommodate pore formation and changes in topology [24, 25], whereas sharp-interface formulations retain the physical free edge and its boundary conditions [26]. A continuum model of post-opening remodeling must therefore couple free-edge mechanics to the dissipation that governs membrane motion. Does hydrodynamic coupling merely change the time scale of post-opening motion, or does it also change the trajectory through membrane-shape space?

Here we study post-opening remodeling using an axisymmetric sharp-interface model of an open lipid membrane in Stokes flow extended from Ref. [27], including free-edge force and moment conditions, bending elasticity, line tension, Gaussian rigidity, spontaneous curvature, local area inextensibility, and membrane viscosity. At the scale of an infected red blood cell, we map the outcomes over line tension and Gaussian rigidity together, including positive Gaussian rigidity, and separately over spontaneous curvature. We track four geometric observables: the edge tangent angle, the neck-to-edge radius ratio, the accumulated rim rotation, and the cap depth. We also formulate and discretize a constrained local-drag gradient flow with the same membrane energy and initial geometry as the hydrodynamic model, so that the effect of the dissipation mechanism can be examined.

After an early edge adjustment from the initial spherical cap, line tension determines whether the pore continues to expand or reverses and shrinks. Positive Gaussian rigidity rotates the rim outward and can produce an internal neck even while the free edge expands. Negative spontaneous curvature instead localizes deformation near the rim, and sufficiently strong curvature causes repeated rolling, with rim rotation preceding cap inversion. Pore radius and membrane energy alone therefore do not describe the remodeling pathway. Edge tangent angle, neck radius, and the order of rim rolling and cap inversion distinguish pathways that reduced descriptions would merge, and the two dissipation models give different pathways in the strongly curved regime. These geometric observables can also be compared with rim curling and membrane eversion during malaria parasite egress.

## Materials and methods

We first summarize the continuum model for an open lipid membrane with a free edge and then introduce two dissipation models for the same membrane energy. Complete governing equations, the discrete schemes, the mobility calibration, and the measurement rules and numerical verification are given in the Supporting Material.

### Open-membrane geometry and energy

We represent the post-opening lipid bilayer as a smooth, orientable open surface Γ(*t*) ⊂ ℝ^3^ whose boundary ∂Γ(*t*) is a free edge that moves with the membrane; lipid flux across the edge is neglected. The bilayer thickness of ∼ 5 nm [28] is small compared with the micrometer-scale vesicles or cells considered here, which supports the two-dimensional surface description standard in continuum membrane mechanics [14, 15]. We impose axisymmetry about the *z* axis and parameterize the meridional generating curve by ***X***(*s, t*) = (*r* (*s, t*), *z*(*s, t*)), where the arclength *s* runs from the pole at *s* = 0 to the free edge at *s* = *l* (*t*) (Fig. 1). The edge is a circle of radius *r*_edge_(*t*) with tangent angle *φ*_edge_(*t*) measured from the horizontal, positive for counterclockwise rotation. We refer to the membrane immediately adjacent to the edge as the “rim.” The unit tangent is ***τ*** = cos *φ* ***e***_*r*_ + sin *φ* ***e***_*z*_, with normal ***n*** = − sin *φ* ***e***_*r*_ + cos *φ* ***e***_*z*_ and outward conormal ***v*** = ***τ*** at the edge. The mean curvature 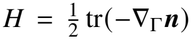 is positive on the initial sphere, the Gaussian curvature is *K* = det(−*∇*_Γ_ ***n***), and the normal and geodesic curvatures of the edge are *κ*_*n*_ = sin *φ*_edge_/*r*_edge_ and *κ*_*g*_ = cos *φ*_edge_/*r*_edge_ (Supporting Material, Table S1).

**Figure 1.**
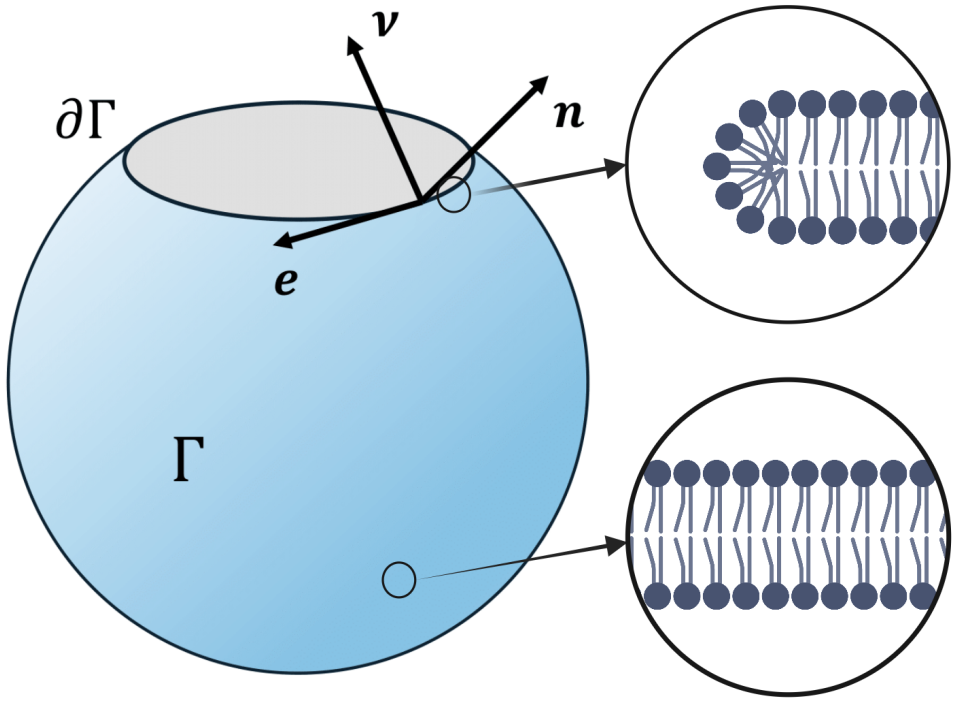
Geometry and notation for an open lipid bilayer. The vectors ***n, v***, and ***e*** denote the surface normal, outward conormal, and edge tangent associated with the membrane Γ and its free edge ∂Γ. The insets illustrate the idealized lipid arrangement within the bilayer and near the free edge.

The elastic driving forces derive from the Helfrich energy of the membrane surface and the line energy of its free edge [14, 16]:

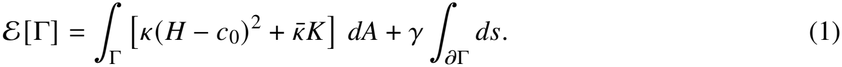

The mean-curvature term penalizes deviations from the spontaneous curvature *c*_0_, with *κ* setting the bending-energy scale. For the disk-topology surfaces considered here, the Gauss–Bonnet theorem transfers the effect of the Gaussian rigidity 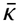 to the force and moment balances at the free edge [16]; for an axisymmetric membrane *∫*_Γ_ *K dA* = 2*π*(1 − cos *φ*_edge_). The line energy, with *γ* the energy per unit edge length, favors a shorter edge [6, 7]. We take *κ, c*_0_, 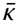, and *γ* to be spatially uniform. Because lipid bilayers have a large area-stretch modulus [29], we impose local area inextensibility [22],

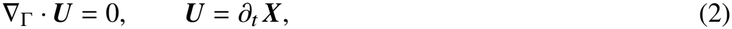

which preserves the area of each material patch while allowing tangential lipid flow. Its Lagrange multiplier *P* is the surface pressure enforcing local area conservation. In the energy convention used here, −*P* is the isotropic area-constraint coefficient, while the complete membrane stress also includes curvature-elastic contributions [30].

With no applied edge force or torque, the free-edge conditions are

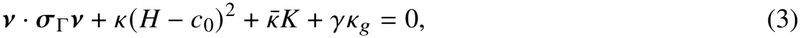

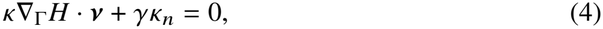

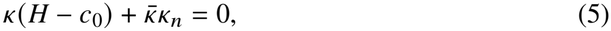

where ***σ***_Γ_ is the surface stress defined below and the geodesic-torsion term vanishes under axisymmetry. The first two equations give the conormal and normal force balances, and the third gives the zero-moment condition; they agree with variational formulations of Helfrich membranes with freely exposed edges [17, 18]. Line tension and Gaussian rigidity therefore enter directly through the free-edge geometry. For a stationary flat disk of radius *R* with *c*_0_ = 0 and no viscous strain, Eq. 3 reduces to −*P* + *γ*/*R* = 0, so the multiplier balances the line force under the area constraint.

### Hydrodynamic coupling

At micrometer scales fluid inertia is negligible [31], so we couple the membrane to an incompressible Newtonian Stokes fluid of viscosity *µ* in Ω = ℝ^3^ \ Γ, with no slip on Γ and decay at infinity. The membrane is a two-dimensional Newtonian fluid with surface viscosity *µ*_Γ_ [21, 22], with stress ***σ***_Γ_ = 2*µ*_Γ_ ***D***_Γ_ (***U***) − *P****P***_Γ_, where ***D***_Γ_ is the in-plane rate-of-deformation tensor and ***P***_Γ_ the tangential projector. The dissipation potential is

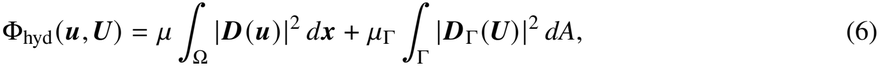

with ***D***(***u***) the bulk rate-of-deformation tensor. We determine the bulk velocity ***u*** and membrane velocity ***U*** from Onsager’s principle, which balances the rate of elastic-energy decrease against the total viscous dissipation subject to bulk incompressibility, no slip, and Eq. 2 [22, 32]. The resulting Stokes equations, surface force balance, and energy law *dℰ*/*dt* = −2Φ_hyd_ are given in Supporting Material, Section S1.

### Local-drag model

To examine the effect of the dissipation mechanism for a fixed membrane energy *ℰ*, we compare the hydrodynamic model with a constrained local-drag *L*^2^ gradient flow driven by that energy. The two models start from the same surface and impose the same local area-inextensibility constraint. The local-drag model omits membrane viscosity and bulk flow, so its *L*^2^ metric has no long-range transmission of forces through the surrounding fluid, and it does not respond to self-contact. Its dissipation potential is

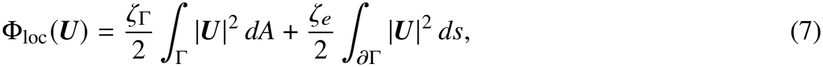

with [*ζ*_Γ_] = N s m^−3^ and [*ζ*_*e*_] = N s m^−2^. Their ratio ℓ_*d*_ = *ζ*_*e*_/*ζ*_Γ_ is a length, and *λ*_*e*_ = ℓ_*d*_/*L* measures the resistance of free-edge motion relative to surface motion. In scaled variables the potential becomes 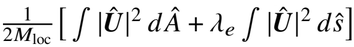 with *M*_loc_ = *κT*/(*ζ*_Γ_ *L*^4^) and the time scale *T* defined below. We fix *λ*_*e*_ = 1 in all local-drag computations. Balancing the decrease of *ℰ* against this dissipation yields the local-drag law [32]. At fixed *λ*_*e*_, varying *M*_loc_ rescales time but leaves the local-drag trajectory in shape space unchanged. The edge drag enters the free-edge force balance and has no counterpart in the hydrodynamic edge conditions.

We integrate the local-drag model with unit mobility in an integration clock 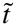 and choose *M*_loc_ so that its energy decrease equals the hydrodynamic decrease at *t*/*T* = 1 for 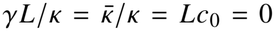. Reported local-drag times are 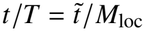. This gives *M*_loc_ = 0.31; over 0 ≤ *t*/*T* ≤ 1 the root-mean-square mismatch of the two energy-decrease histories is 0.20 *κ*, or 6.6% of the matched decrease. The fitted *M*_loc_ is held fixed in all comparisons.

### Numerical methods

We discretize the hydrodynamic model with an axisymmetric boundary-element/finite-element method [27]. A Stokes single-layer potential represents the bulk flow, and a mixed finite-element formulation on the generating curve couples the fluid response to the surface force balance, local area inextensibility, and membrane motion, using an inf–sup-stable *P*_2_–*P*_1_ velocity–pressure pair, auxiliary curvature fields, and a semi-implicit backward-Euler step. Because the traction-jump density behaves as *d*^−1/2^ with distance *d* from the free edge, the mesh is graded by a sine map that concentrates cells there. The logarithmic singularity of the single-layer kernel is integrated with a hybrid Gauss–trapezoidal rule [33]. The local-drag model is discretized with a semi-implicit parametric finite-element method using piecewise-linear positions and curvature vectors and piecewise-constant pressures on an arclength-uniform mesh with *N* = 64 cells; a candidate step is accepted only if the discrete energy is nonincreasing. Both discrete schemes are given in full in Supporting Material, Section S2.

Across the simulations, relative area and lipid-mass conservation hold to within 10^−3^ in the line-tension and Gaussian-rigidity sweeps and within 1.6% in the strongly curved rolling regime; numerical verification and stopping criteria are summarized in Supporting Material, Section S4.

### Parameters

The simulations are organized by the dimensionless groups

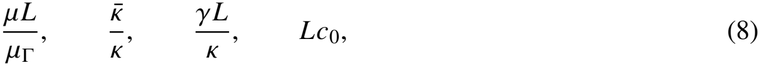

which compare bulk-fluid with membrane-surface dissipation, Gaussian-curvature and line energies with the bending-energy scale, and the membrane size with the spontaneous-curvature length |*c*_0_|^−1^. Balancing the bending-energy release rate *κ*/*T* with surface-viscous dissipation *µ*_Γ_ *L*^2^/*T* ^2^ gives the characteristic time *T* = *L*^2^*µ*_Γ_/*κ*, and time is reported as *t*/*T*. We take *L* = 3 *µ*m, comparable to the radius of an approximately spherical infected red blood cell immediately before egress [1, 2], and use *κ* = 100 *k*_*B*_*T* as a reference bending-energy coefficient in the convention of Eq. 1. In the alternative convention *E*_*b*_ = (*k*_*b*_/2) ∫_Γ_ (2*H* − *C*_0_)^2^ *dA*, matching the normal orientation gives *κ* = 2*k*_*b*_ and *c*_0_ = *C*_0_/2. Reference viscosities are *µ* = 10^−3^ Pa s in the bulk and *µ*_Γ_ = 10^−7^ N s m^−1^ on the membrane, a surface-viscosity scale between reported values for lipid bilayers and red-cell membranes [2, 34, 35]. Table 1 summarizes the reference values and sampled ranges; for the reference values these ranges correspond to *γ* = 0–0.17 pN, *c*_0_ = −1.67 to 0 *µ*m^−1^, and 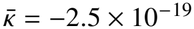 to 1.6 × 10^−19^ J.

**Table 1.** Dimensional reference values and their mapping to the computational model. These values set the simulation scales, and 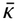 is varied over positive and negative values.

| Quantity | Units | Baseline | Mapping or sweep |
|---|---|---|---|
| *L* | m | 3 × 10<sup>−6</sup> | length scale |
| *μ* | Pa s | 10<sup>−3</sup> | *μL*/*μ*<sub>Γ</sub> = 0.03 |
| *μ*<sub>Γ</sub> | N s m<sup>−1</sup> | 10<sup>−7</sup> | *T* = 2.17 s |
| *κ* | J | 100 *k*<sub>*B*</sub>*T* | bending rigidity |
| *κ̄* | J | varied | *κ̄*/*κ* ∈ [−0.6, 0.4] |
| *γ* | N | varied | *γL*/*κ* ∈ [0.4, 1.2] (line-tension sweep), [0, 1.2] (map) |
| *c*<sub>0</sub> | m<sup>−1</sup> | varied | *Lc*<sub>0</sub> ∈ [−5, 0] |

Both solvers start from the same spherical cap with radius *L* = 3 *µ*m and initial pore radius *r*_0_ = 0.5 *µ*m, interpolated onto their respective finite-element spaces, and all computations are carried out in scaled variables. The hydrodynamic runs use *N* = 64 cells unless stated otherwise; the *Lc*_0_ = −5 run of Fig. 5 uses *N* = 128. The local-drag integration step is 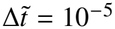 with unit mobility.

## Results

### Line tension selects pore expansion, long-lived opening, or shrinkage

To isolate the effect of line tension, we set 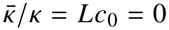 and vary *γL*/*κ*. Comparing a planar disk with a closed sphere at the initial area *A*_0_ gives the disk–sphere energy crossover at 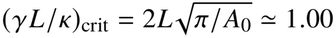 (Supporting Material, Section S5), and we sample line tensions on both sides of this value.

All pores initially expand, including those that later reverse and shrink (Fig. 2A). The initial spherical cap has *H* = 1/*L* at the free edge, while the zero-spontaneous-curvature moment condition requires *H* = 0. The resulting curvature mismatch is relaxed through an early deformation localized near the rim (Fig. 2B). The initial increase in pore radius is therefore a short-time curvature-relaxation effect rather than the long-time response to line tension.

**Figure 2.**
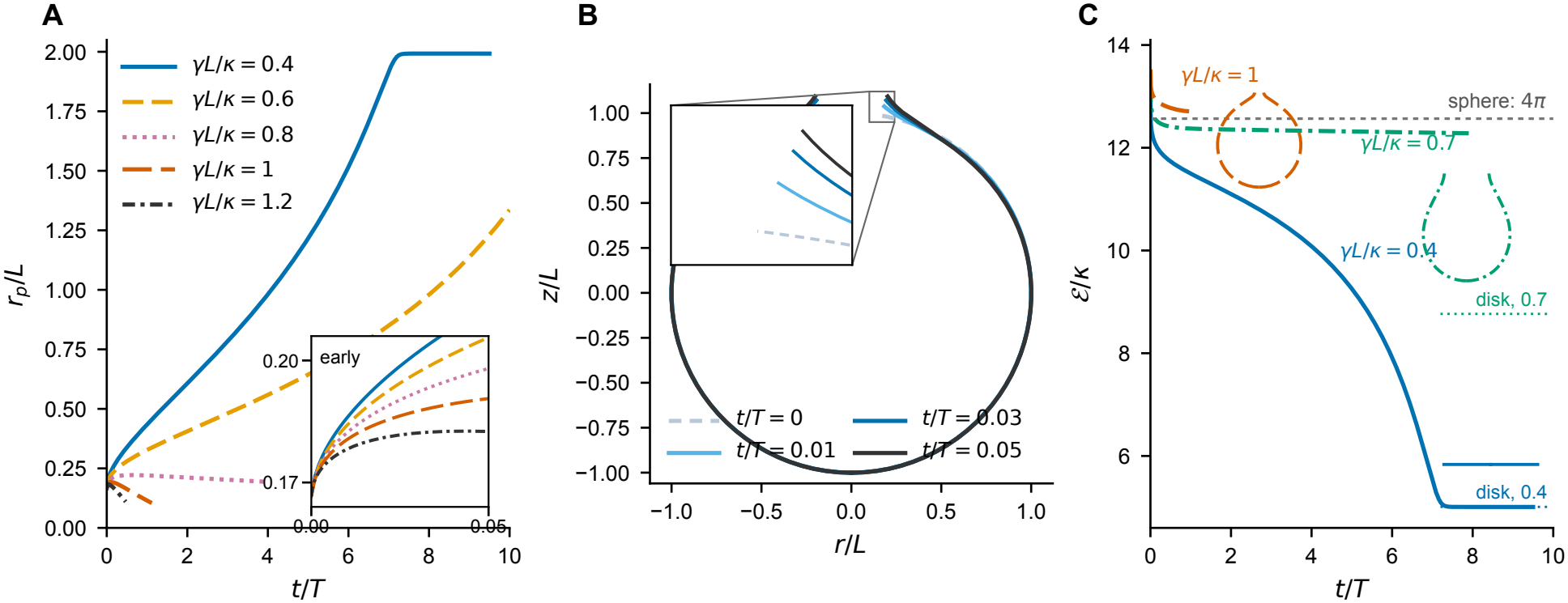
Membrane evolution as line tension is varied. (A) Pore radius for *γL*/*κ* = 0.4, 0.6, 0.8, 1.0, and 1.2, with 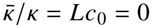. (B) Early profiles for *γL*/*κ* = 0.7 at *t*/*T* = 0, 0.01, 0.03, and 0.05, showing that the initial deformation is concentrated near the rim. (C) Scaled membrane energy E/*κ* for *γL*/*κ* = 0.4, 0.7, and 1.0; insets show the final computed profile for each case.

After the initial transient, the trajectories separate into three regimes. For *γL*/*κ* = 0.4 and 0.6, the pore continues to expand, whereas for *γL*/*κ* = 0.8, 1.0, and 1.2 it reverses and shrinks (Fig. 2A). At the intermediate value *γL*/*κ* = 0.7, the membrane remains in a long-lived intermediate configuration: the energy decays slowly while a finite pore persists, with *r*_edge_/*L* ≃ 0.48 at *t*/*T* = 8, the end of the simulated interval (Fig. 2C). Finite-time reversal is bracketed between *γL*/*κ* = 0.6 and 0.8, below the disk–sphere energy crossover.

In ruptured lipid vesicles, variation in the effective line tension could contribute to large differences in pore lifetime. For *L* = 3 *µ*m and *κ* = 100 *k*_*B*_*T*, the long-lived case *γL*/*κ* = 0.7 corresponds to *γ* ≃ 0.10 pN, roughly two orders of magnitude below typical values reported for bare lipid pores [6]. Membrane additives can reduce line tension by a similar factor and extend pore lifetimes from seconds to minutes [6]. Edge-binding proteins can have related effects; for example, talin localizes at membrane free edges and stabilizes open, cup-shaped liposomes [5].

### Gaussian rigidity changes rim orientation and promotes internal neck formation

We next set *Lc*_0_ = 0 and vary the Gaussian rigidity. The Gaussian-curvature energy 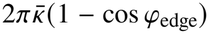 depends on the rim orientation, while the zero-moment condition, Eq. 5, links 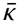 to the local rim curvature. With *k*_*s*_ and *k* _*θ*_ the meridional and azimuthal curvatures at the circular edge, *H* = (*k*_*s*_ + *k* _*θ*_)/2 and *κ*_*n*_ = *k* _*θ*_, so the moment condition gives

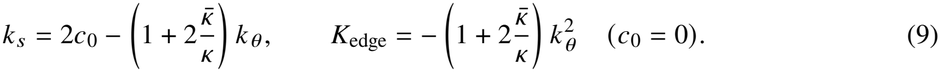

For *k* _*θ*_ ≠ 0 the edge is therefore saddle-like for every 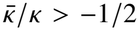, including 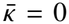, and *K*_edge_ = 0 when *k* _*θ*_ = 0. Positive 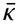 increases the magnitude of the opposing meridional curvature required for a given *k* _*θ*_ . Whether this edge condition produces an internal neck depends on how *k* _*θ*_ evolves, which the simulations determine. Line tension acts differently: the line energy is proportional to the edge length and hence to *r*_edge_.

We fix *γL*/*κ* = 0.5 and compare 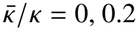, 0.2, and 0.4 (Fig. 3). By *t*/*T* = 2, larger Gaussian rigidity has rotated the rim farther outward: 1 − cos *φ*_edge_ decreases from 0.83 to 0.39 and 0.12 across the three cases (Fig. 3A), and *r*_neck_/*r*_edge_ decreases from 0.99 to 0.83 and 0.58 (Fig. 3B). For 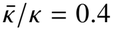, the neck criterion *r*_neck_/*r*_edge_ < 0.8 is first reached at *t*/*T* ≃ 0.40, and the constriction develops inside the rim, behind the free edge (Fig. 3C). Between *t* = 0 and *t*/*T* = 2, *r*_edge_/*L* increases from 0.17 to 0.30, so the edge moves outward while the internal neck narrows. The neck-to-edge radius ratio measures this constriction and distinguishes it from the saddle geometry at the rim itself.

**Figure 3.**
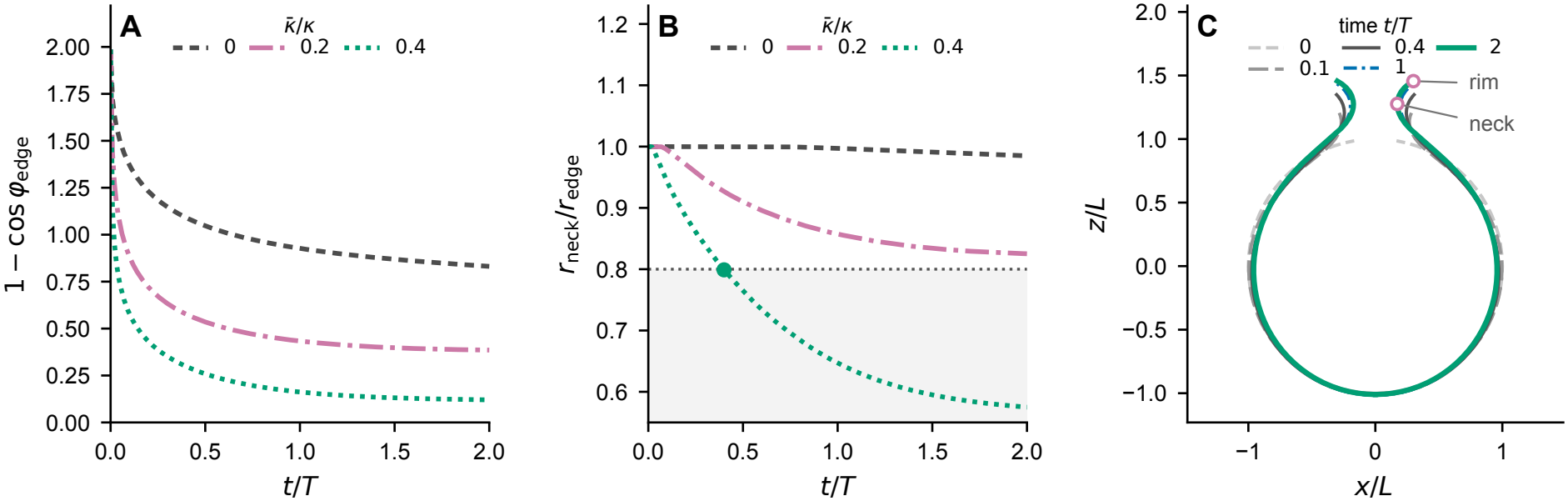
Positive Gaussian rigidity promotes formation of an internal neck. (A) Rim orientation, measured by 1 − cos *φ*_edge_, and (B) neck-to-edge radius ratio for 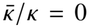, 0.2, and 0.4, with *γL*/*κ* = 0.5 and *Lc*_0_ = 0. In (B), the ratio is set to 1 until an interior minimum in radius develops; the detection rule is given in Supporting Material, Section S4. The dotted line marks the neck criterion, and the green point its first crossing. (C) Meridional profiles for 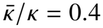; magenta circles mark the neck and free edge in the final profile.

To place this behavior in a broader parameter range, we vary Gaussian rigidity and line tension together over 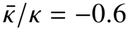 to 0.4 and *γL*/*κ* = 0 to 1.2 in increments of 0.2 (Supporting Material, Section S5, Figs. S1 and S2). We classify a trajectory as neck-forming if *r*_neck_/*r*_edge_ < 0.8 at any output time. Neck formation occurs at 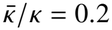 and 0.4 for every line tension in the map. For 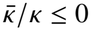, increasing *γL*/*κ* instead changes the outcome from expansion to direct shrinkage, and every case in this range shrinks when *γL*/*κ* ≥ 0.8.

In equilibrium calculations with prescribed pores, Gaussian rigidity alters the membrane shape near the edge [19]. In the present dynamics, an outward-moving edge can also coexist with a narrowing internal neck, so the edge tangent angle and neck radius, rather than pore radius alone, distinguish uniform expansion from localized constriction. Gaussian moduli of ordinary lipid bilayers are generally negative [36]; the positive values used here test how the sign of 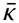 changes the free-edge dynamics.

### Negative spontaneous curvature localizes rim rolling, with implications for malaria parasite egress

To isolate the effect of spontaneous curvature, we set 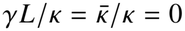 and vary *Lc*_0_. After unwrapping the edge tangent angle *φ*_edge_(*t*) in time, we define the accumulated rim rotation by

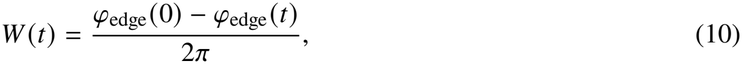

so that *W* = 1 corresponds to one full turn of the rim. We characterize the cap by the signed depth *h* = *z*_rim_ − *z*_pole_, where *z*_rim_ is the first maximum of *z* on the branch from the outermost-radius point to the free edge, and we say that the cap inverts when *h* becomes negative, the model counterpart of the cap eversion observed during egress (Supporting Material, Section S4). For *Lc*_0_ = 0 and −1, the rim reaches only *W* = 0.47 and 0.84, respectively, by *t*/*T* = 5 (Fig. 4A and B). The first complete turn within the simulated interval occurs at *Lc*_0_ = −1.5, with *W* = 1 at *t*/*T* = 2.32. Stronger negative spontaneous curvature produces sustained, repeated rolling: for *Lc*_0_ = −5, the rim completes its first turn by *t*/*T* = 0.11 and reaches three turns by *t*/*T* = 1.17.

**Figure 4.**
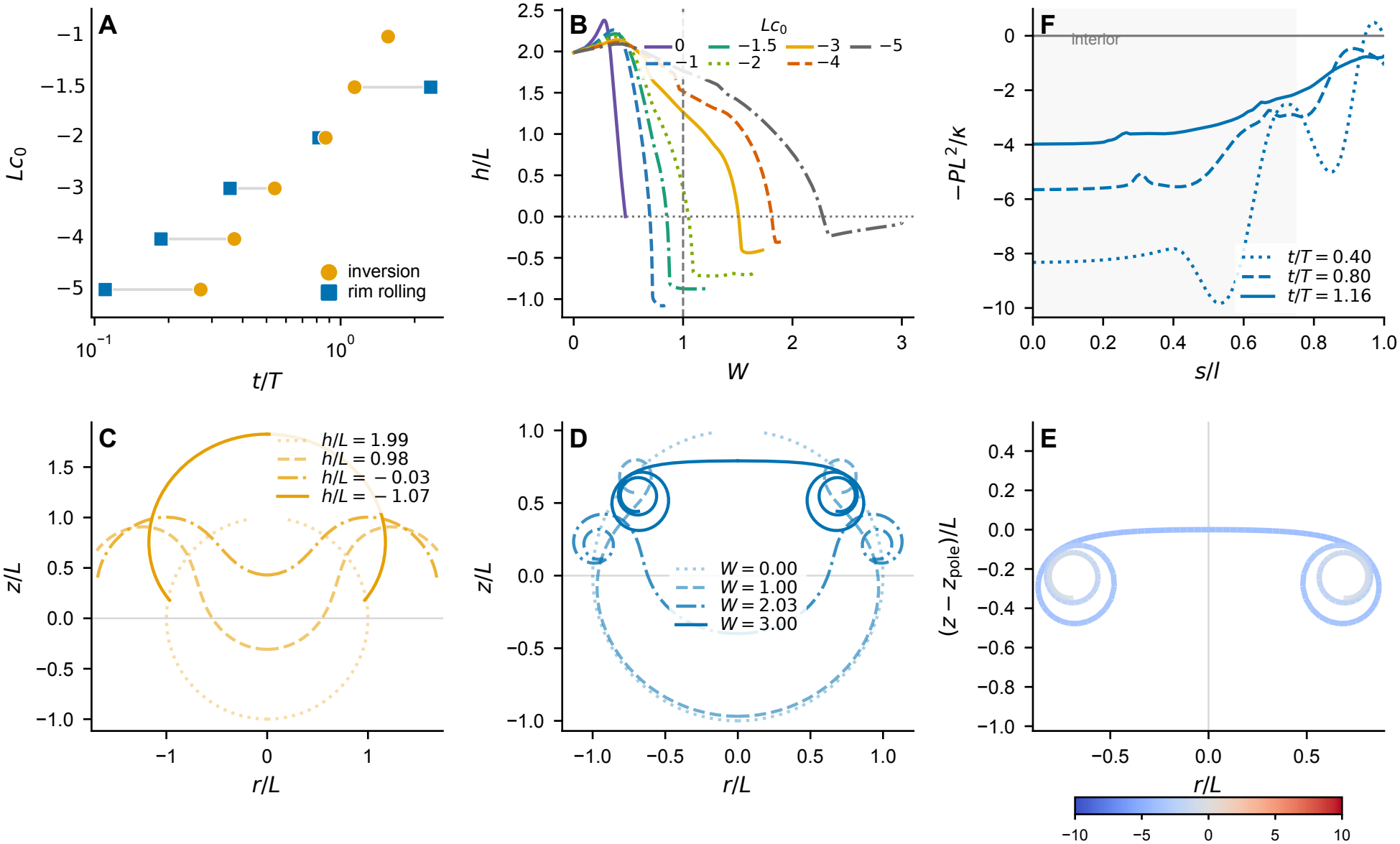
Negative spontaneous curvature drives cap inversion and localized rim rolling. (A) First-passage times for cap inversion (orange circles) and one complete rim turn (blue squares). Lines connect the two events for each value of *Lc*_0_. No complete rim turn is reached at *Lc*_0_ = 0 (*W* = 0.47) or −1 (*W* = 0.84) within *t*/*T* ≤ 5, and the cap does not invert at *Lc*_0_ = 0; these events are not plotted. The computations at *Lc*_0_ = −2, −3, and −4 stop when neighboring membrane branches reach the proximity cutoff, and the *Lc*_0_ = −5 computation terminates at *t*/*T* = 1.17 (Supporting Material, Section S4). (B) Signed cap depth versus accumulated rim rotation for *Lc*_0_ = 0, −1, −1.5, −2, −3, −4, and −5. Reference lines mark *h* = 0 and one full rim turn, *W* = 1. (C) Mirrored profiles for *Lc*_0_ = −1 at *h*/*L* = 1.99, 0.98, −0.03, and −1.07, corresponding to *t*/*T* = 0, 1.35, 1.56, and 5. (D) Mirrored profiles for *Lc*_0_ = −5 at *W* = 0, 1.00, 2.03, and 3.00, corresponding to *t*/*T* = 0, 0.11, 0.23, and 1.17. (E) Area-constraint multiplier −*PL*^2^/*κ* for *Lc*_0_ = −5 at *t*/*T* = 1.16 and *W* = 3.00; blue indicates negative values. (F) Profiles of −*PL*^2^/*κ*, in the Helfrich-energy convention of Eq. 1, at (*t*/*T, W*) = (0.40, 2.61), (0.80, 2.91), and (1.16, 3.00) for *Lc*_0_ = −5. Shading marks the pole-side region *s*/*l* ≤ 0.75.

The location of the deformation also changes with *Lc*_0_ (Fig. 4C and D). At *Lc*_0_ = −1, most of the shape change is distributed over the cap and the rim does not complete a full turn by *t*/*T* = 5. At *Lc*_0_ = −5, the cap initially changes little while repeated rolling is confined to a narrow region near the free edge. This localization reflects the free-edge moment condition. With zero line tension and Gaussian rigidity, the edge must satisfy *H* = *c*_0_, whereas the initial spherical cap has *H* = 1/*L*, so the curvature mismatch grows as *Lc*_0_ becomes more negative. The spontaneous-curvature length |*c*_0_|^−1^ = *L*/|*Lc*_0_| also decreases, which favors a more localized response and shifts the motion from broad cap deformation toward rim rolling.

Strong rolling produces large spatial variations in the area-constraint multiplier (Fig. 4E and F); this variation, unlike the sign and mean of the multiplier, does not depend on the energy convention. In the Helfrich-energy convention of Eq. 1, at *t*/*T* = 1.16 (*W* = 3.00), −*PL*^2^/*κ* is negative throughout the pole-side region *s*/*l* ≤ 0.75, where its area-weighted mean is −3.28. Removing the constant 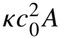 from the energy leaves the motion unchanged and shifts this mean by (*Lc*_0_)^2^ to 21.72.

During malaria parasite egress, the infected red blood cell membrane forms a pronounced outward-rolled rim before the remaining cap inverts [1]. Callan-Jones et al. proposed parasite-induced spontaneous curvature as a mechanism for this rolling [2]. In the present simulations, the cap inverts before the first complete rim turn at *Lc*_0_ = −1.5, whereas the first complete turn precedes cap inversion for *Lc*_0_ ≤ −2 (*c*_0_ ≤ −0.67 *µ*m^−1^; Fig. 4A). The latter ordering matches the observed egress sequence qualitatively.

### Dissipation model changes rolling at strong spontaneous curvature

Spontaneous curvature supplies the driving energy for the rolling described above, but the path taken through shape space also depends on the dissipation mechanism. To examine this effect, we compare the hydrodynamic and local-drag models using the same membrane energy and initial shape, with the local-drag mobility calibrated from the matched energy decay at *Lc*_0_ = 0 (Methods). After this time rescaling, the two *Lc*_0_ = 0 trajectories pass through similar cap-like shapes over 0 ≤ *t*/*T* ≤ 1 (Fig. 5A). Their root-mean-square contour distance is 0.07*L* at *t*/*T* = 0.25 and 0.11*L* at *t*/*T* = 1, while the final edge radii differ by only 0.03*L*. At zero spontaneous curvature, a single rescaling of time captures most of the difference between the two dissipation models.

**Figure 5.**
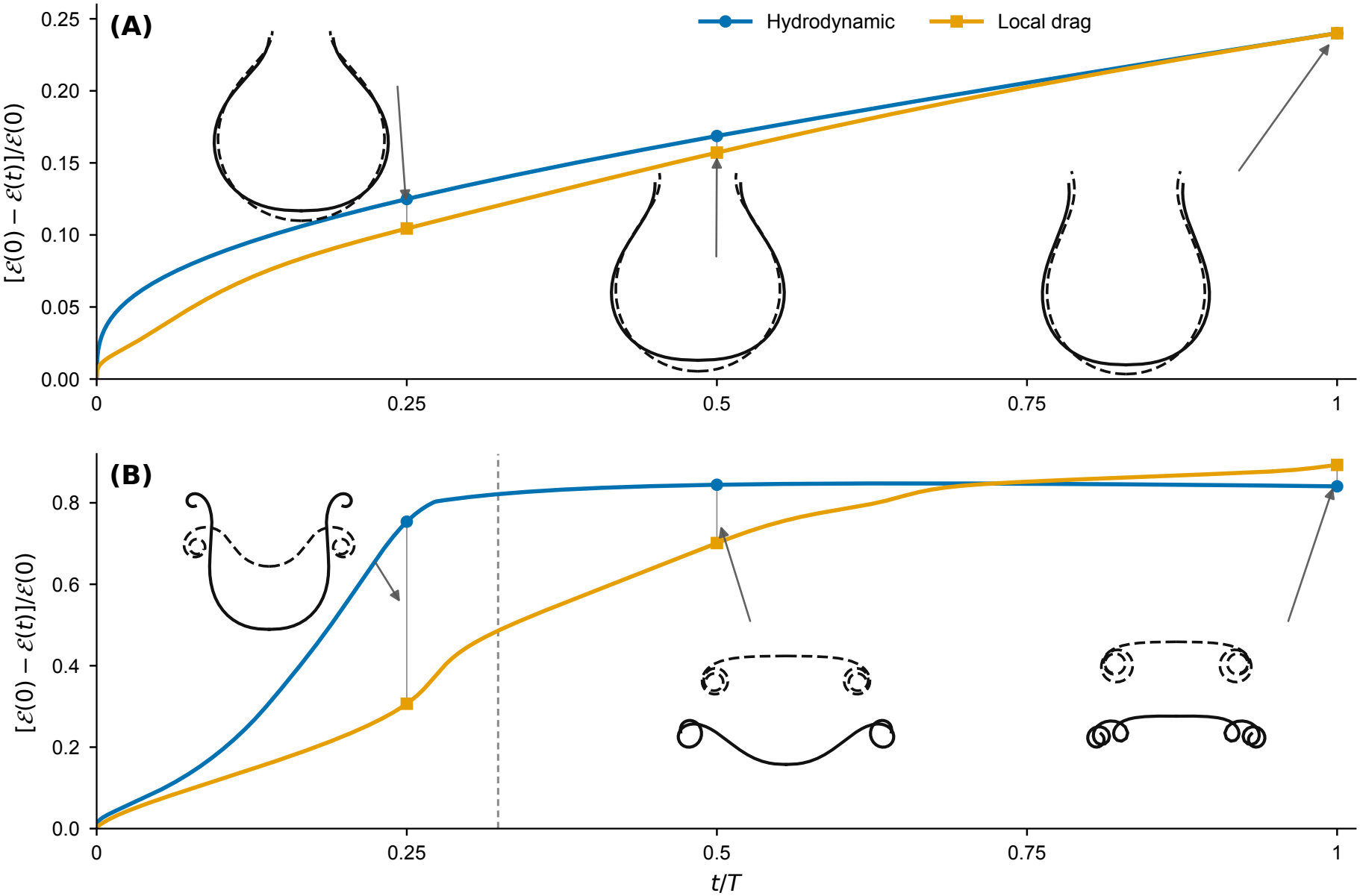
Hydrodynamic and local-drag trajectories follow similar shapes at zero spontaneous curvature but diverge during strong rolling. (A) Fractional energy decrease, [*ℰ*(0) − *ℰ*(*t*)]/*ℰ*(0), and membrane profiles at *t*/*T* = 0.25, 0.5, and 1 for *Lc*_0_ = 0. Contour distances are unweighted root-mean-square distances over 1024 arclength-uniform points at equal arclength fraction from pole to rim. The vertical offset of the profiles is for display only. (B) Energy decrease and profiles for *Lc*_0_ = −5. The local-drag profile first self-intersects between the output times *t*/*T* = 0.29 and 0.32; the vertical gray line marks *t*/*T* = 0.32, the first output state with an intersection. Blue circles and dashed black profiles denote the hydrodynamic model, while orange squares and solid black profiles denote the local-drag model. Arrows connect profiles to the corresponding energy values. The last two hydrodynamic profiles in (B) are shifted upward for clarity. Both panels use 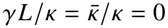, the same initial cap, and the same scaled coordinate limits.

A single rescaling does not reconcile the two models in the strongly curved regime. At *Lc*_0_ = −5, the local-drag model retains a deeper cap and localizes more deformation near the rim than the hydrodynamic model (Fig. 5B; hydrodynamic *N* = 128, local-drag *N* = 64). The contour distance reaches 0.70*L* by *t*/*T* = 0.25, about ten times the corresponding value at *Lc*_0_ = 0. By *t*/*T* = 1, the hydrodynamic and local-drag models have dissipated 84% and 89%, respectively, of the initial energy *ℰ*(0) defined by Eq. 1. The local-drag profile has by then crossed itself, whereas the hydrodynamic profile does not self-intersect over the computed interval. Comparable energy decay therefore does not imply comparable membrane motion once strong rolling develops.

One plausible contributor to this difference is hydrodynamic resistance in the narrow gaps created by rolling. As neighboring membrane branches approach, fluid must drain from the gap, generating lubrication pressure that opposes further normal motion. Similar drainage-controlled slowing has been found for nearby vesicle membranes [37], consistent with classical lubrication results for deformable drops and rigid bodies [38, 39].

## Discussion

After pore formation, the free-edge moment balance sets the local rim geometry, while the dissipation model influences the trajectory through shape space, as the comparison at *Lc*_0_ = −5 shows.

Line tension becomes important only after the initial edge-relaxation transient, during which every pore first expands as the spherical-cap rim adjusts toward the free-edge moment condition. Weak line tension then allows continued expansion, whereas stronger line tension reverses the edge motion and drives shrinkage, and the case *γL*/*κ* = 0.7 retains a finite opening throughout the simulated interval. In experiments, membrane additives and edge-binding proteins can reduce the effective line tension and substantially prolong pore lifetime [5, 6].

Gaussian rigidity acts through the free-edge force and moment balances and can change the local geometry without reversing the direction of edge motion: for positive 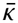, an internal neck develops behind an edge that is still expanding, so an increasing pore radius does not imply uniform opening of the entire pore region. This distinction is absent from reduced models based only on pore radius, membrane tension, and line tension [6, 7, 12, 13].

Negative spontaneous curvature changes both where deformation occurs and the order of geometric events. As *Lc*_0_ decreases, the spontaneous-curvature length becomes shorter and the deformation localizes progressively closer to the free edge. At *Lc*_0_ = −1, cap inversion occurs while a complete rim turn is not reached within the simulated interval. At *Lc*_0_ = −1.5, cap inversion precedes the first complete turn, whereas the first turn occurs earlier for *Lc*_0_ ≤ −2. The ordering at sufficiently negative spontaneous curvature is relevant to malaria parasite egress, where a rolled rim forms before the remaining red-cell membrane everts [1], and is consistent with the mechanism proposed in Ref. [2].

The comparison of the two dissipation models shows that membrane energy alone does not determine the rolling trajectory. At *Lc*_0_ = 0, where the local-drag time scale is calibrated, the two models pass through similar cap-like shapes. At *Lc*_0_ = −5, both have dissipated about 85–90% of the initial energy by *t*/*T* = 1 but reach qualitatively different geometries: the local-drag profile self-intersects, whereas the hydrodynamic profile remains separated over the computed interval. This separation is consistent with lubrication resistance as neighboring layers approach [37–39]. Isolating its contribution would require a matched-resolution comparison with near-contact quadrature in which only the hydrodynamic coupling changes. A related mechanism may help keep rolled membrane layers separated during malaria egress.

The model has several limitations. We consider passive axisymmetric motion only after a pore has formed, treat cytoskeletal remodeling implicitly, take spontaneous curvature to be homogeneous, and omit pore nucleation, active parasite forces, spatial heterogeneity in membrane composition, azimuthal buckling, and explicit self-contact. The erythrocyte spectrin network is strongly remodeled and rapidly collapses near parasite egress [40, 41], motivating the effective passive-surface approximation at this stage. For *Lc*_0_ = −5, we use *c*_0_ = −1.67 *µ*m^−1^, whose magnitude is about one quarter of the estimate of Callan-Jones et al. in the convention used here [2]. Their estimate is 13 ± 5 *µ*m^−1^ in the (*k*_*b*_/2)(2*H* − *C*_0_)^2^ convention and 6.5 ± 2.5 *µ*m^−1^ in the convention of Eq. 1. Finite simulation horizons and proximity cutoffs also limit conclusions about complete pore closure and late near-contact dynamics. Extending the model to three dimensions, heterogeneous curvature, and explicit contact will be necessary to determine whether the strongly rolled state is stable to non-axisymmetric perturbations and how rolled membrane layers evolve after close approach.

The most direct experimental test of these results would track pore radius, edge tangent angle, neck radius, and cap depth in the same rupture movie. Such measurements would show whether rim rolling precedes cap inversion and whether an expanding pore coexists with an internal constriction. A series of vesicles with additives that lower the line tension [6] would test the predicted change from shrinkage through a long-lived opening to continued expansion as *γL*/*κ* decreases.

## Data and code availability

Code and data are available at https://github.com/han-zhou-math/free-edge-membrane-dynamics.

## Author contributions

H.Z., Y.-W.C., and Y.M. designed the research and developed the model. H.Z. developed the numerical methods, wrote the software, performed the simulations, and wrote the manuscript. Y.-W.C. and Y.M. reviewed and edited the manuscript.

## Acknowledgments

Y.-W.C. was supported by a National Institutes of Health R35 Award (R35GM156396) and a Burroughs Wellcome Fund Investigators in the Pathogenesis of Infectious Disease Program Award (1022785). Y.M. was partially supported by the National Science Foundation through the University of Pennsylvania Materials Research Science and Engineering Center (MRSEC) (DMR-2309043).

## Supporting material

Supporting Material is available with this article. It contains the complete governing equations and orientation conventions (Section S1), the discrete boundary-element/finite-element and local-drag schemes (Section S2), the parameter mapping and mobility calibration (Section S3), the measurement rules and numerical verification (Section S4), and the extended Gaussian-rigidity sweeps and pathway map (Section S5, Figs. S1 and S2).

## Declaration of generative AI and AI-assisted technologies in the manuscript preparation process

During the preparation of this work the authors used Claude (Anthropic) and Codex (OpenAI) in order to assist with writing simulation, analysis, and figure-generation code and to improve the language and readability of the manuscript. After using these tools, the authors reviewed and edited the content as needed and take full responsibility for the content of the publication.

## Supporting Material

This Supporting Material accompanies the article *Free-edge mechanics and hydrodynamic coupling shape membrane remodeling after pore opening*. Section S1 gives the complete governing equations and orientation conventions, Section S2 the discrete schemes, Section S3 the parameter mapping and mobility calibration, Section S4 the measurement rules and numerical verification, and Section S5 the extended Gaussian-rigidity sweeps and pathway map. Equations, figures, and tables carry an S prefix; “main-text Fig. *n*” and “main-text Eq. *n*” refer to the article. All lengths, times, and energies are scaled by *L, T* = *L*^2^*µ*_Γ_*/κ*, and *κ* unless units are stated.

### S1 Governing equations and orientation conventions

We impose axisymmetry about the *z* axis and parameterize the meridional generating curve by ***X***(*s, t*) = (*r*(*s, t*), *z*(*s, t*)), where *s* is arclength. The curve runs from the symmetry axis at *s* = 0 to the free edge at *s* = *l*(*t*). The edge is a circle of radius *r*_edge_(*t*) with tangent angle *φ*_edge_(*t*) measured from the horizontal, taking clockwise rotation as negative and counterclockwise rotation as positive. Increasing arclength runs from the pole to the free edge, with unit tangent ***τ*** = cos *φ* ***e***_*r*_ + sin *φ* ***e***_*z*_, normal ***n*** = sin *φ* ***e***_*r*_ + cos *φ* ***e***_*z*_, and outward conormal ***ν*** = ***τ*** at *s* = *l*(*t*). In this orientation the initial spherical cap has *H* = +1*/L*. The normal and geodesic curvatures of the edge are *κ*_*n*_ = −sin *φ*_edge_*/r*_edge_ and *κ*_*g*_ = + cos *φ*_edge_*/r*_edge_, and axisymmetry gives zero geodesic torsion, *τ*_*g*_ = 0. The curvature tensor, mean curvature, and Gaussian curvature are

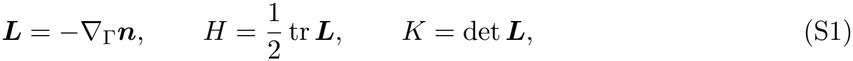

where *∇*_Γ_ is the surface gradient. The mean curvature *H* is the signed average of the principal curvatures, while *K* is their product. The chosen orientation fixes the signs of *H, c*_0_, *κ*_*n*_, and *κ*_*g*_ throughout; Table S1 summarizes these conventions. With this definition of *κ*_*g*_ the Gauss–Bonnet theorem Jfor the disk-topology surface reads ∫_Γ_ *K dA* + ∫_*∂*Γ_ *κ*_*g*_ *ds* = 2*π*, which gives the axisymmetric Identity ∫_Γ_ *K dA* = 2*π*(1 − cos *φ*_edge_) used in the main text.

**Table S1:**
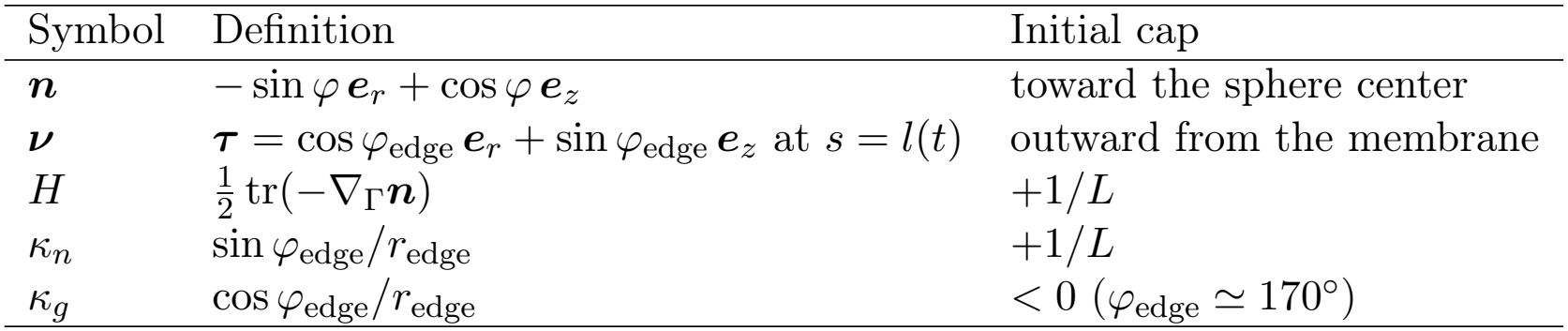
Orientation conventions used throughout, with their values on the initial spherical cap of radius *L* and pore radius *r*_0_ = *L/*6.

| Symbol | Definition | Initial cap |
|---|---|---|
| $\boldsymbol{n}$ | $-\sin\varphi\,\boldsymbol{e}_r+\cos\varphi\,\boldsymbol{e}_z$ | toward the sphere center |
| $\boldsymbol{\nu}$ | $\boldsymbol{\tau}=\cos\varphi_{\mathrm{edge}}\,\boldsymbol{e}_r+\sin\varphi_{\mathrm{edge}}\,\boldsymbol{e}_z$ at $s=l(t)$ | outward from the membrane |
| $H$ | $\frac{1}{2}\,\mathrm{tr}(-\nabla_{\Gamma}\boldsymbol{n})$ | $+1/L$ |
| $\kappa_n$ | $\sin\varphi_{\mathrm{edge}}/r_{\mathrm{edge}}$ | $+1/L$ |
| $\kappa_g$ | $\cos\varphi_{\mathrm{edge}}/r_{\mathrm{edge}}$ | $<0$ ($\varphi_{\mathrm{edge}}\simeq 170^{\circ}$) |

The elastic driving forces derive from the Helfrich energy of the membrane surface and the line energy of its free edge [1, 2]:

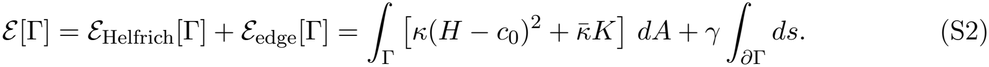

The Helfrich energy contains mean-curvature and Gaussian-curvature contributions. The mean-curvature term penalizes deviations from the spontaneous curvature *c*_0_; *κ* sets the bending-energy scale and *H* = *c*_0_ is the locally preferred mean curvature. For the disk-topology surfaces considered here, the Gauss–Bonnet theorem transfers the effect of the Gaussian rigidity 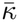 to the natural force and moment balances at the free edge [2]. The final term is the line energy, with *γ* denoting the energy per unit edge length and favoring a shorter edge [3, 4]. We take *κ, c*_0_, 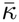, and *γ* to be spatially uniform.

Because lipid bilayers have a large area-stretch modulus [5], area changes are small on the deformation scales considered here. We therefore impose local area inextensibility, as is standard in continuum models of fluid membranes [6]:

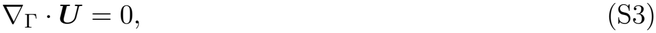

where ***U*** = *∂*_*t*_***X***. This constraint preserves the area of each material patch while allowing tangential lipid flow and changes in membrane shape. Its Lagrange multiplier *P* is the surface pressure enforcing local area conservation. In the energy convention used here, −*P* is the isotropic area-constraint coefficient. The complete membrane stress also includes curvature-elastic contributions [7].

#### S1.1 Local-drag dissipation

The local-drag model replaces the hydrodynamic dissipation by the potential

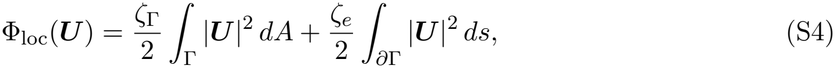

with [*ζ*_Γ_] = N s m^*−*3^ and [*ζ*_*e*_] = N s m^*−*2^, subject to the same local area-inextensibility constraint, Eq. S3, whose multiplier contributes − ∫_Γ_ *P ∇*_Γ_ · ***V*** *dA* to the weak velocity equation. Balancing the decrease of *ℰ* against this dissipation yields the local-drag law; its scaled form is given in Section S3.

#### S1.2 Hydrodynamic coupling

The membrane is coupled to an incompressible Newtonian Stokes fluid of bulk viscosity *µ* in Ω = ℝ^3^ \Γ, with no slip on Γ and decay at infinity, and is itself a two-dimensional Newtonian fluid with surface viscosity *µ*_Γ_. For the bulk velocity ***u*** and membrane velocity ***U***, the rate-of-deformation tensors are

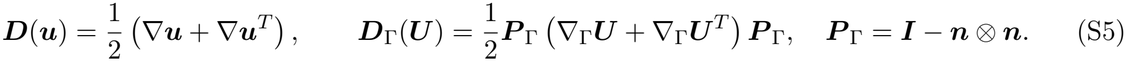

Here ***D*** is the bulk rate-of-deformation tensor, while the projected tensor ***D***_Γ_ measures the corresponding in-plane membrane deformation. The dissipation potential is

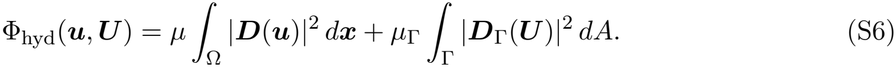

The first term accounts for viscous dissipation in the surrounding fluid and the second for viscous dissipation within the membrane. We determine ***u*** and ***U*** from Onsager’s principle, which balances the rate of elastic-energy decrease against total viscous dissipation [6, 8]. The admissible velocities satisfy

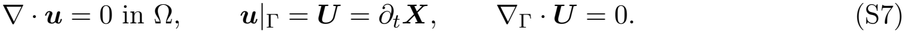

These conditions enforce bulk incompressibility, no slip between fluid and membrane, and local membrane-area conservation, respectively. The bulk pressure *p* and surface pressure *P* act as Lagrange multipliers for the two divergence constraints. The corresponding stresses are

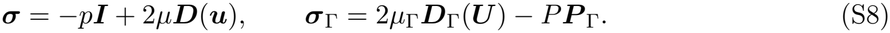

The pressure terms enforce incompressibility, while the viscous terms resist deformation. The force-balance equations are

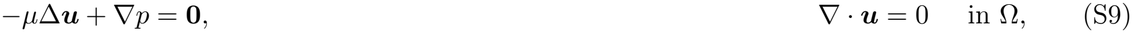

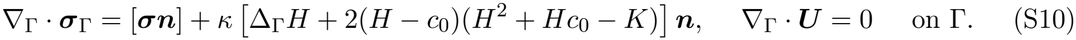

In the bulk, pressure and viscous stress satisfy the Stokes equations. On Γ, the divergence of the surface stress balances the bulk traction jump [***σn***] and the normal Helfrich force. For spatially uniform 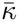, the Gaussian-curvature energy contributes to the free-edge conditions but produces no interior membrane force [2].

With no applied edge force or torque, the free-edge conditions are

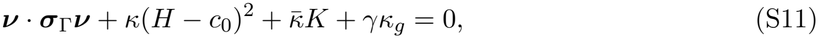

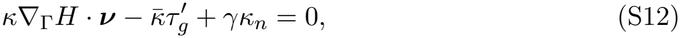

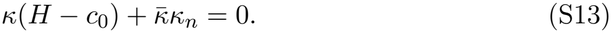

The first two equations give the conormal and normal force balances, and the third gives the zero-moment condition. Line tension and Gaussian rigidity therefore enter directly through the free-edge geometry. These boundary conditions agree with variational formulations of Helfrich membranes with freely exposed edges [9, 10]. A prime denotes differentiation with respect to edge arclength, and *τ*_*g*_ = 0 under axisymmetry. For a stationary flat disk of radius *R* with *c*_0_ = 0 and no viscous strain, Eq. S11 reduces to −*P* + *γ/R* = 0, so the multiplier balances the line force under the area constraint.

The coupled system obeys the energy law

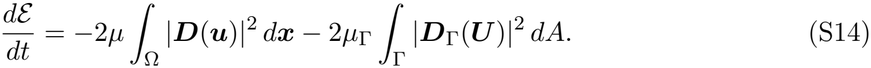

Hence the membrane energy decreases through bulk and surface viscous dissipation.

For the circular edge, 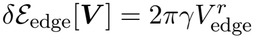. With *κ*_*g*_ = cos *φ*_edge_*/r*_edge_ and *κ*_*n*_ = sin *φ*_edge_*/r*_edge_, this variation per unit edge length is *γκ*_*g*_***V*** · ***ν*** − *γκ*_*n*_***V*** · ***n*** = (*γ/r*_edge_)*V* ^*r*^, and the corresponding line force is −*γκ*_*g*_***ν*** + *γκ*_*n*_***n*** = −(*γ/r*_edge_)***e***_*r*_. The conormal part of this force enters the discrete force balance of Section S2 through the term 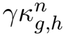 and the normal part through the boundary term 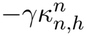 of the bending-force equation, consistent with the strong conditions in Eqs. S11–S13.

### S2 Discrete schemes

We discretize the hydrodynamic model with an axisymmetric boundary-element/finite-element (BEM–FEM) method [11]. A boundary-integral representation accounts for viscous flow in the surrounding fluid, while a mixed finite-element formulation on the generating curve couples the fluid response to surface force balance, local area inextensibility, and membrane motion.

#### S2.1 BEM–FEM scheme for the hydrodynamic model

##### Axisymmetric boundary-integral reduction

For axisymmetric flow with no azimuthal component, we represent the bulk velocity by the Stokes single-layer potential

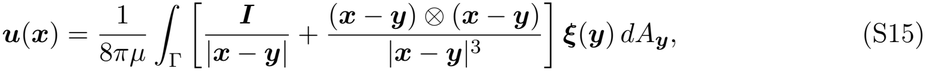

where ***ξ*** = [***σn***] is the unknown traction-jump density. Analytical integration over the azimuthal angle reduces Eq. S15 to a one-dimensional operator *S*[***ξ***] on the generating curve, with a logarithmic singularity on the diagonal.

Let *t*^*n*^ = *n*Δ*t*, and let 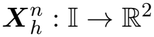, with I = (0, 1), parameterize the discrete generating curve 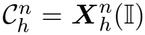. A field *f* on 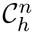 is represented in the parameter coordinate by 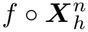. For scalar or vector fields on 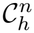, we use the axisymmetric weighted products

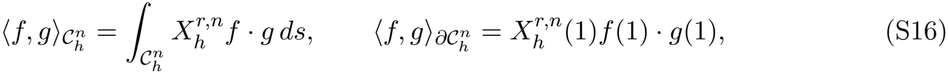

where the common factor 2*π* from azimuthal integration is omitted. A subscript *s* denotes differentiation with respect to arclength on 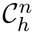, and superscripts *r* and *z* denote cylindrical components.

We partition I into cells *I*_*i*_ = [*α*_*i−*1_, *α*_*i*_] and define

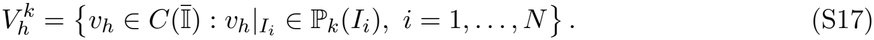

The vector space

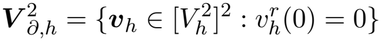

enforces the axis condition at *r* = 0. We also define

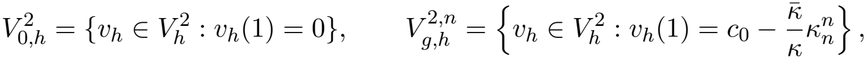

where the affine constraint in 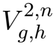 imposes the discrete zero-moment condition at the free edge.

Near the free edge, the traction-jump density behaves as |***ξ***| ∼*d*^*−*1*/*2^, where *d* is the distance to the edge. To resolve this singular behavior, we use the graded mesh

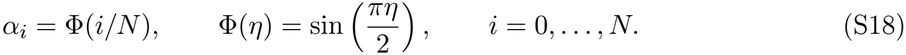

This grading produces cells of size *O*(*N* ^*−*2^) near the free edge and *O*(*N* ^*−*1^) in the interior, concentrating resolution where the traction density is singular.

##### Mixed finite-element scheme

To avoid higher spatial derivatives in the finite-element formulation, we introduce an auxiliary mean-curvature vector ***H***, the scalar mean curvature *H*, and a normal bending-force density *g*. Given the discrete generating curve 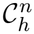, we seek

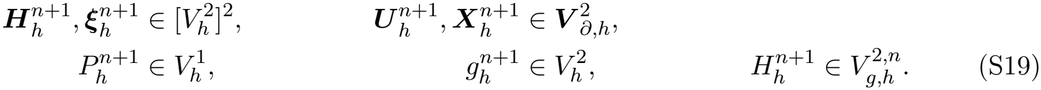

For the axisymmetric surface Stokes problem, the velocity–pressure pair 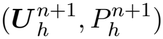 is discretized with the inf–sup-stable *P*_2_–*P*_1_ Taylor–Hood pair. The geometry, traction-jump density, and curvature auxiliary fields are also represented by continuous quadratic elements.

Let 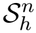 denote the discrete axisymmetric single-layer operator. A superscript *h* indicates Gauss– Legendre quadrature, whereas *h, \** denotes the quadrature used for the double integral involving 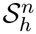. Time is discretized with a semi-implicit backward-Euler scheme. The highest-order and linear terms are treated implicitly to improve stability:

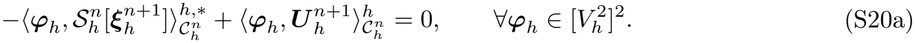

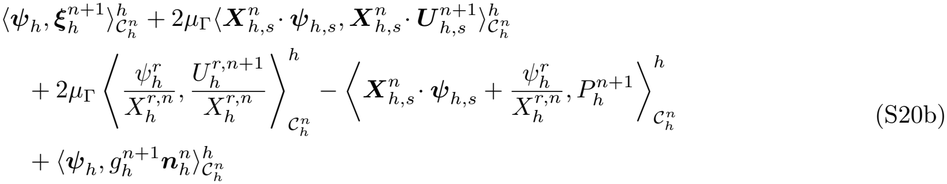

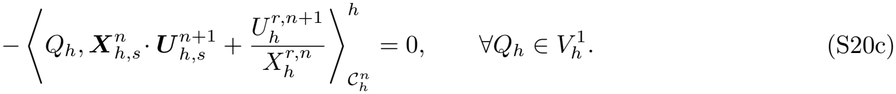

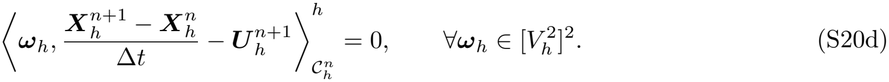

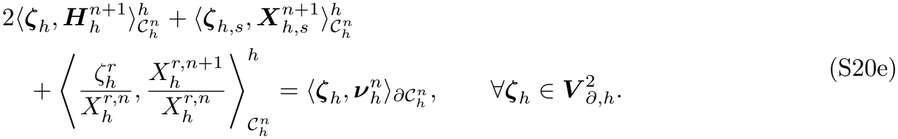

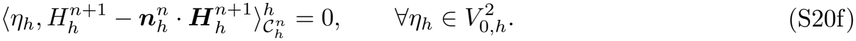

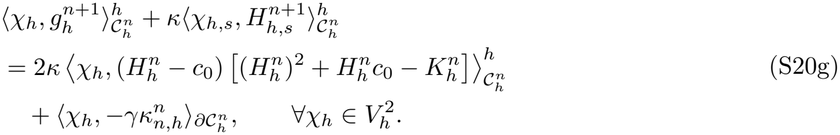

Equations S20a–S20g give, respectively, the boundary-integral velocity relation, surface force balance, local area-inextensibility constraint, kinematic update, weak curvature-vector identity, scalar curvature relation, and normal bending-force equation. The boundary term proportional to 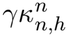 and the affine constraint defining 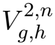 enforce the normal-force and zero-moment conditions at the free edge.

The coefficients 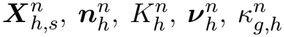, and 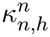 are evaluated on the known curve 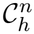. The resulting linear system simultaneously determines the position, velocity, traction-jump density, pressure, and auxiliary curvature fields at *t*^*n*+1^. The highest-order terms 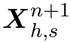 and 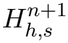 are implicit, whereas nonlinear geometric coefficients and free-edge data are lagged to *t*^*n*^. The matrix is sparse apart from the dense block associated with 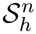.

##### Quadrature

Regular finite-element integrals are evaluated with five-point Gauss–Legendre quadrature. For the logarithmically singular double integral in Eq. S20a, we use a hybrid Gauss–trapezoidal rule [12].

#### S2.2 FEM for the local-drag model

We discretize the constrained local-drag gradient flow with a semi-implicit parametric finite-element method. Within the scaled local-drag equations the symbols *κ*, 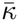, *γ*, and *c*_0_ stand for 1, 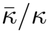, *γL/κ*, and *Lc*_0_. We retain the time levels, parameter interval, and geometric notation introduced above, but use continuous piecewise-linear elements for the position and meridional curvature vector and piecewise constants for the surface pressure. On the local-drag mesh, let

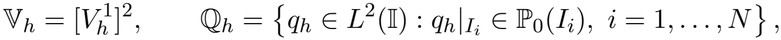

where 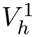 is defined on this partition. During each time step, *ds*^*n*^, 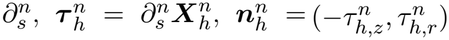, and 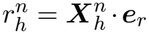 are evaluated on 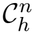 and held fixed; 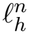 denotes the current meridional length. Given 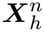 and 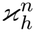, we solve for

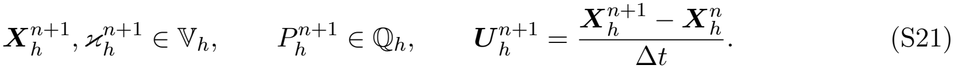

Here 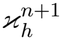 denotes the meridional curvature vector and 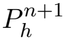 enforces local area inextensibility.

After omitting the common azimuthal factor 2*π*, the axisymmetric energy takes the form 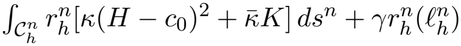. We approximate its first variation by setting

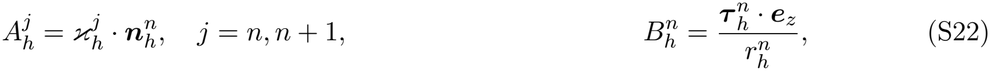

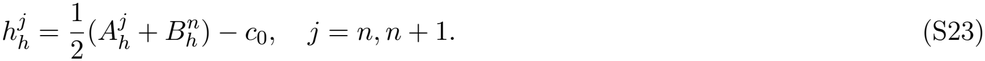

At the pole, the quotient defining 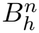 is replaced by its regular limit 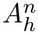. A linear update is obtained by using

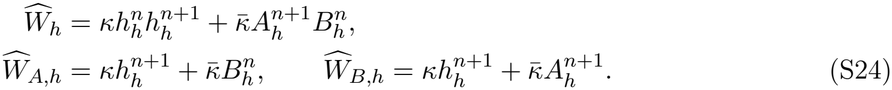

The weak energy variation involves the three coefficients

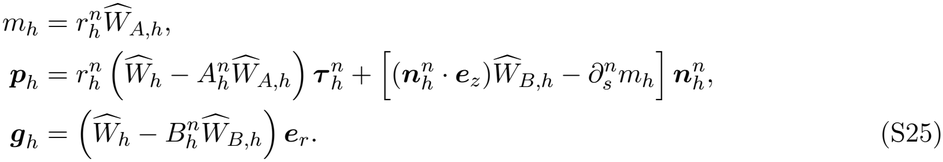

The derivative of *m*_*h*_ is evaluated cellwise. All three quantities are linear in 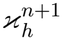 and carry the scaled coefficients 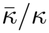 and *Lc*_0_.

The fully discrete scheme is summarized below. We use 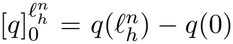 and the weighted products from Eq. S16. The unknowns in Eq. S21 satisfy the following equations for all **Ψ**_*h*_ ∈ V_*h*_, *χ*_*h*_ ∈ Q_*h*_, and ***V*** _*h*_ ∈ V_*h*_ with 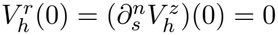:

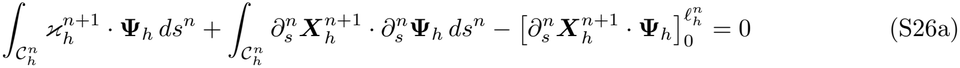

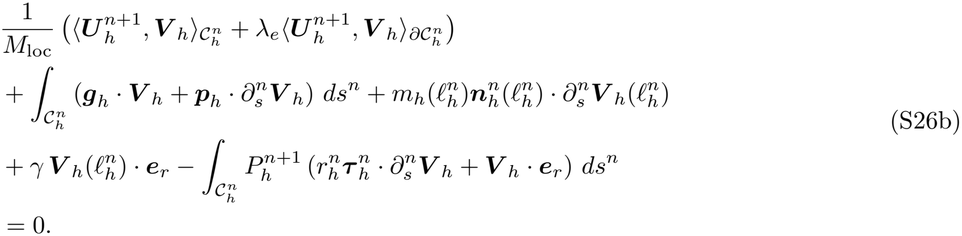

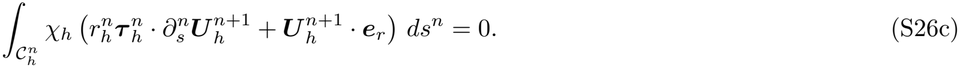

The pole conditions are imposed strongly and are stated separately from the weak equations:

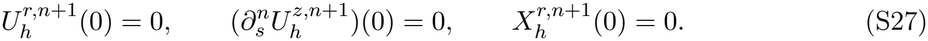

The kinematic update preserves the last condition, which is reset after each accepted step to eliminate roundoff drift. The pole is an endpoint of the generating curve but not a physical boundary of the membrane. Because 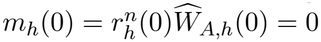, the explicit energy boundary term vanishes there and appears only at the free edge in Eq. S26b. We retain the endpoint terms in Eq. S26a because they define the meridional curvature vector weakly without imposing an artificial condition on 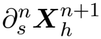.

Equation S26b combines local surface and edge drag with the bending, Gaussian-curvature, linetension, and pressure terms. Equation S26c enforces local area inextensibility cell by cell. Because the metric and geometric coefficients are evaluated at *t*^*n*^, each step requires the solution of a linear saddle-point system. Regular finite-element integrals are evaluated with five-point Gauss–Legendre quadrature. Step-acceptance rules are given in Section S4.

### S3 Parameters, scaling, and mobility calibration

The dimensional parameters have units 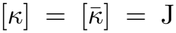, [*γ*] = N, [*c*_0_] = m^*−*1^, [*µ*] = Pa s, and [*µ*_Γ_] = N s m^*−*1^. Length is scaled by *L* and energy by *κ*. A membrane velocity of order *L/T* corresponds to a surface strain rate of order *T* ^*−*1^; balancing the bending-energy release rate *κ/T* with surface-viscous dissipation *µ*_Γ_*L*^2^*/T* ^2^ gives *T* = *L*^2^*µ*_Γ_*/κ*, which is 2.17 s for the reference values *L* = 3 *µ*m, *µ*_Γ_ = 10^*−*7^ N s m^*−*1^, and *κ* = 100 *k*_*B*_*T* . With 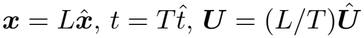, and energy scale *κ*, the local-drag dissipation potential becomes

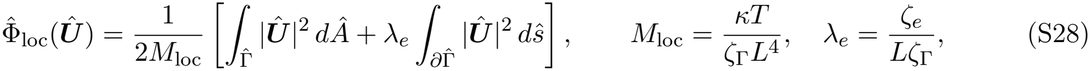

so that 1*/M*_loc_ is the overall dimensionless resistance and *λ*_*e*_ the relative edge resistance. We fix *λ*_*e*_ = 1 in all local-drag computations.

Both solvers start from the same spherical cap with radius *L* and initial pore radius *r*_0_ = *L/*6 (0.5 *µ*m), interpolated onto their respective finite-element spaces. The local-drag trajectories use an arclength-uniform mesh with *N* = 64 and fixed mesh topology, whereas the hydrodynamic solver uses the sine-graded mesh of Eq. S18. The local-drag integration step is 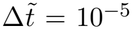 in the unit-mobility clock 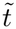.

We calibrate *M*_loc_ by integrating the local-drag model with unit mobility and choosing *M*_loc_ so that its energy decrease equals the hydrodynamic decrease at *t/T* = 1 for 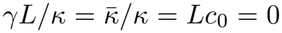; both histories are interpolated linearly. Reported local-drag times are 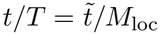. The procedure gives *M*_loc_ = 0.31. Over 0 *t/T* 1 the root-mean-square mismatch of the two energy-decrease histories is 0.20 *κ*, or 6.6% of the matched decrease, and the maximum mismatch is 0.42 *κ*. This value is held fixed in all comparisons of main-text Fig. 5. For the reference values, *M*_loc_ = 0.31 corresponds to *ζ*_Γ_ = 3.6 × 10^4^ N s m^*−*3^ and *ζ*_*e*_ = 1.1 × 10^*−*1^ N s m^*−*2^.

For the reference values the sampled ranges *γL/κ* ∈ [0, 1.2], *Lc*_0_ ∈ [−5, 0], and 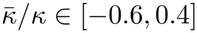 correspond to *γ* = 0–0.17 pN, *c*_0_ = −1.67 to 0 *µ*m^*−*1^, and 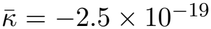 to 1.6 × 10^*−*19^ J. The long-lived opening at *γL/κ* = 0.7 corresponds to *γ* ≃ 0.10 pN.

### S4 Measurement rules and numerical verification

In the line-tension and Gaussian-rigidity sweeps, relative area and lipid-mass errors remain below 10^*−*3^, positive energy excursions are at most 10^*−*3^|*ℰ*(0)|, and the discrete energy–dissipation residual 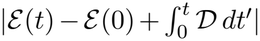 is within 5% of the energy drop, where D = −*dℰ /dt* is the dissipation rate in Eq. S14. In the spontaneous-curvature sweep (main-text Fig. 4), the maximum relative area errors are 1.5 × 10^*−*4^, 1.1 × 10^*−*3^, 2.0 × 10^*−*3^, 3.1 × 10^*−*3^, 7.6 × 10^*−*3^, 1.6 × 10^*−*2^, and 1.6 × 10^*−*2^ for *Lc*_0_ = 0, − 1, − 1.5, − 2, − 3, − 4, and − 5.

The local-drag solver enforces monotone energy decrease by step halving and resolving the system (up to 24 halvings, floor 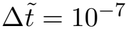, energy tolerances 10^*−*12^), uses a linear-solver tolerance of 10^*−*8^, limits the node step to 0.25 of the cell length and the minimum cell to 0.05 of the mean, and stops if the relative area drift exceeds 2%. The local-drag solver monitors the cellwise divergence residual (≤5 × 10^*−*9^). Runs terminate at the maximum simulated time or upon reaching a closure, opening, or interbranch-proximity threshold.

The neck radius *r*_neck_ is the interior minimum of *r* sought on the rim-side 15% of arclength of the profile resampled on 4096 arclength-uniform points, using a two-sided stencil of 0.01 in arclength so that neither the pole nor the free edge can be selected; the ratio *r*_neck_*/r*_edge_ is set to 1 until such a minimum develops. We classify a profile as having an internal neck when *r*_neck_*/r*_edge_ *<* 0.8. Rim-rotation milestones (*W* = 0.5, 1) are linearly interpolated between saved states. Cap inversion is the first of three consecutive saved states with *h <* −10^*−*3^*L*, taken at saved-frame resolution. The edge tangent angle is unwrapped along arclength and then in time. The rim height *z*_rim_ is the first maximum of *z* on the branch from the outermost-radius point to the free edge.

For the spontaneous-curvature family, halving the time step from 2 × 10^*−*4^ to 10^*−*4^ changes the first-passage times for cap inversion and for one complete rim turn by at most 3.6 × 10^*−*3^ in *t/T* (with a saved-frame resolution of 0.01 for inversion), whereas the separation between the two events is at least 0.05 for *Lc*_0_ = −1.5 to −4; for each *Lc*_0_, the separation exceeds the refinement change for the same *Lc*_0_ by a factor of about 18 or more. For *Lc*_0_ = −5, *r*_edge_*/L* differs by 6.1 × 10^*−*3^ between *N* = 64 and *N* = 128 over 0 ≤ *t/T* ≤ 0.145. Ten Gaussian-rigidity cases at *γL/κ* ∈ {0, 1} have time-step pairs (10^*−*3^ and 2.5 × 10^*−*4^) whose edge radii differ by at most 7.9 × 10^*−*3^*L*.

### S5 Extended Gaussian-rigidity sweeps and pathway map

A useful energetic reference is obtained by comparing a planar disk and a closed sphere at fixed area. For area *A*_0_, the disk has energy 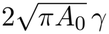, whereas the sphere has energy 4*πκ*. Equating the two gives

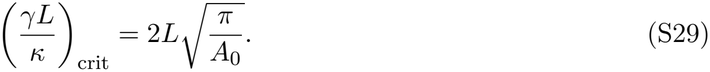

For the initial cap, *A*_0_*/*(*πL*^2^) ≃ 3.97, so (*γL/κ*)_crit_ ≃ 1.00. We therefore sample line tensions on both sides of this reference value.

To place this behavior in a broader parameter range, we vary Gaussian rigidity and line tension together. At fixed area, the Gaussian term changes the closed-sphere energy to 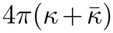 but leaves the planar-disk energy unchanged. Their energetic crossover therefore occurs at

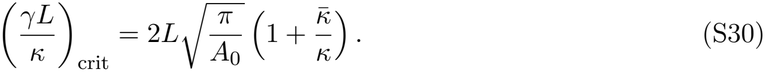

We compare the outcomes of the two-parameter sweep with this reference curve.

The edge-radius histories in Fig. S1 show a systematic change with line tension. With *γL/κ* = 0, the edge expands for every sampled 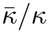 (Fig. S1D). At *γL/κ* = 0.5, negative-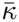 cases shrink, whereas the zero and positive cases move outward (Fig. S1E). At *γL/κ* = 1, cases with nonpositive 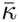 shrink, while positive 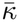 produces an initial expansion and a finite edge radius over the displayed interval (Fig. S1F). Thus, among the sampled cases with nonzero line tension, negative Gaussian rigidity is associated with direct pore shrinkage, whereas positive Gaussian rigidity supports outward rim rotation and neck formation.

**Figure S1:**
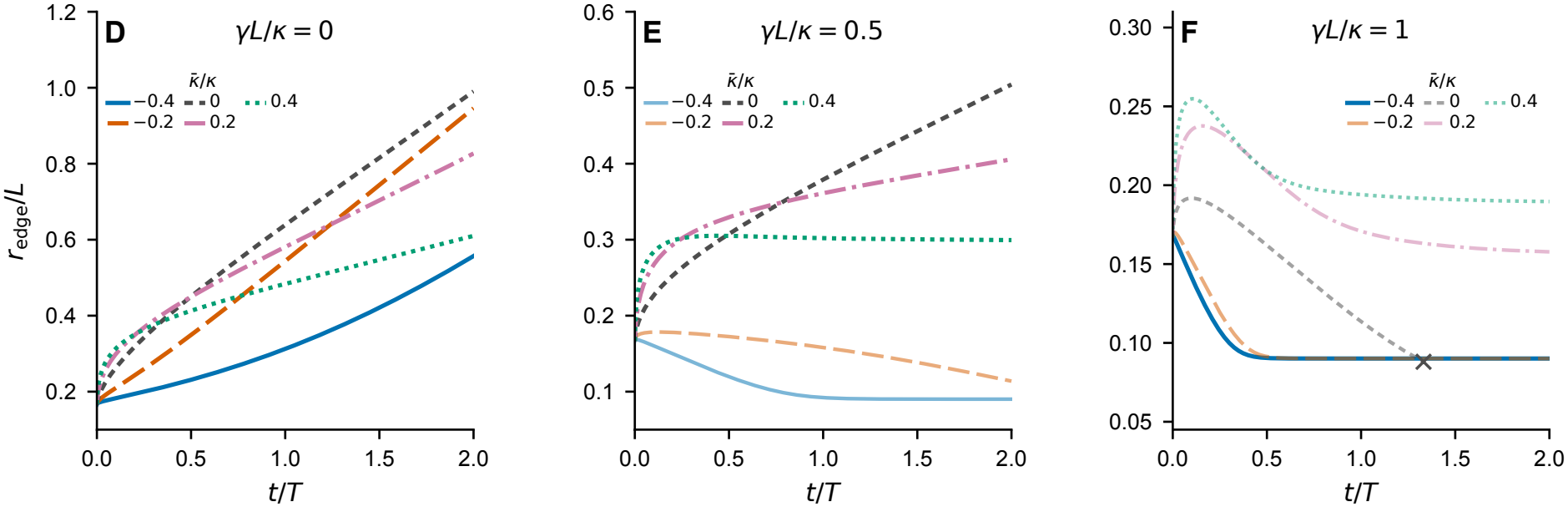
Edge-radius histories across Gaussian rigidity and line tension. (D–F) Edge radius for 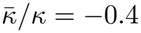 to 0.4 at *γL/κ* = 0, 0.5, and 1, with *Lc*_0_ = 0. Panel letters continue those of main-text Fig. 3. The cross in (F) marks the termination point of the 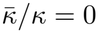 trajectory.

Because an outward-moving edge can coexist with an internal constriction, pore radius alone is not sufficient to classify the morphology. We label a trajectory as neck-forming if *r*_neck_*/r*_edge_ *<* 0.8 at any saved time. For the pathway map the interior minimum is sought on the rim-side 45% of arclength with the same resampling and stencil as in Section S4. Among trajectories without a neck, a net change larger than 2% in *r*_edge_*/L* distinguishes expansion from shrinkage; for smaller changes, we use the late-time trend together with the accumulated edge displacement.

The resulting pathway map summarizes the 42 sampled outcomes on the 0.2 parameter grid (Fig. S2). Neck formation occurs at the sampled positive values 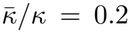 and 0.4 across the investigated line tensions. For 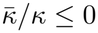, increasing *γL/κ* instead drives a transition from expansion to direct shrinkage, and every sampled case in this range shrinks when *γL/κ* ≥ 0.8. The dashed curve is the disk–sphere energetic crossover, and the symbols are simulation outcomes.

**Figure S2:**
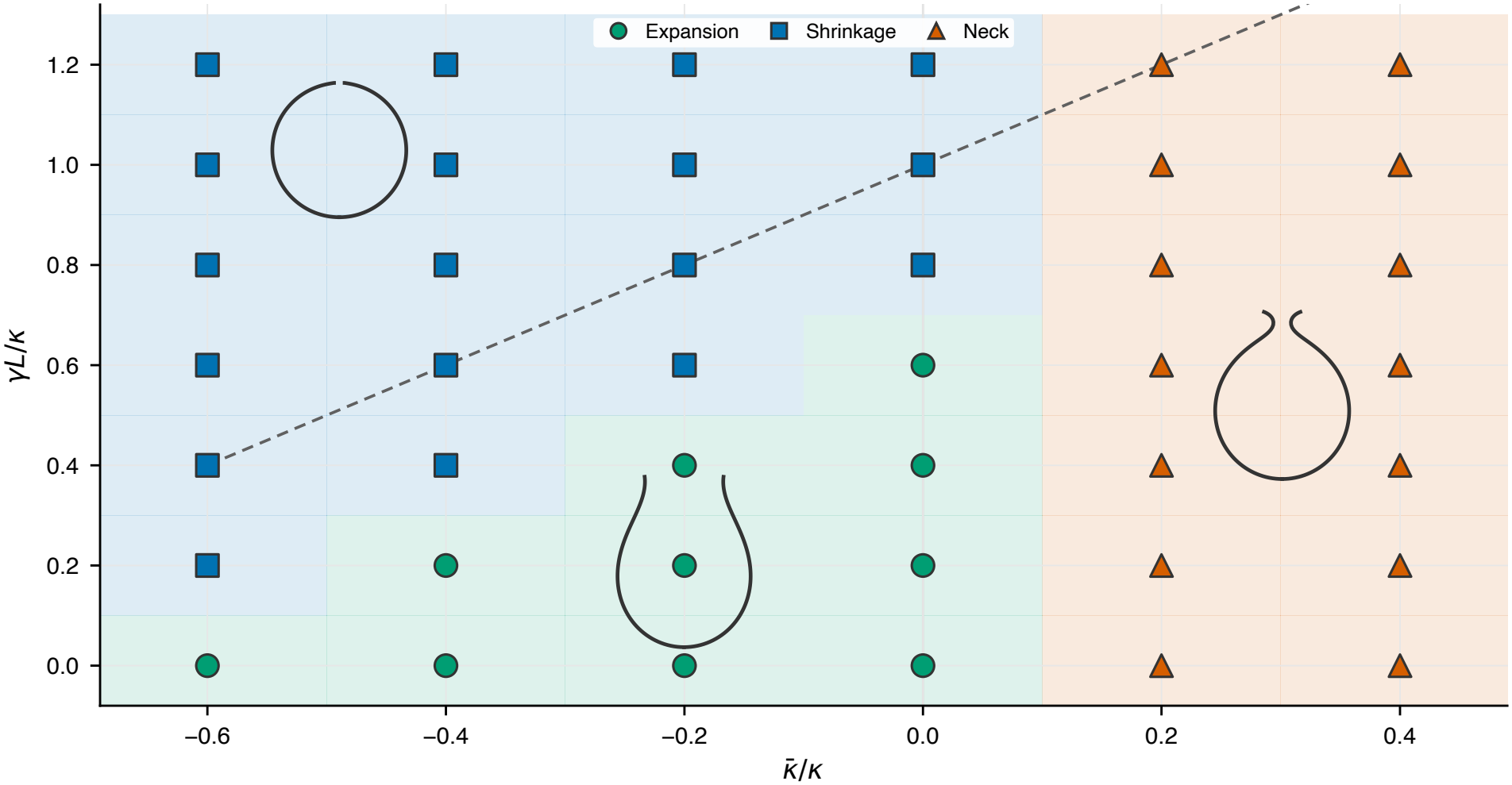
Membrane pathways across line tension and Gaussian rigidity. The map samples 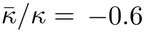 to 0.4 and *γL/κ* = 0 to 1.2 in increments of 0.2. Circles denote expansion, squares direct shrinkage, and triangles neck formation. A trajectory is classified as neck-forming if *r*_neck_*/r*_edge_ *<* 0.8 at any saved time; edge displacement and its late-time trend classify the remaining cases as expanding or shrinking. Colored cells display the discrete outcomes without interpolation. The dashed line shows the disk–sphere energetic crossover from Eq. S30. Insets show representative final profiles and are positioned for legibility.

